# TGF-β signalling regulates the balance between protective and regulatory CD4⁺ T cell responses in visceral leishmaniasis

**DOI:** 10.64898/2026.08.03.742418

**Authors:** Jinrui Na, Fabian de Labastida Rivera, Teija M. Frame, Luzia Bukali, Jessica A. Engel, Christian R. Engwerda

## Abstract

Visceral leishmaniasis (VL) is a potentially fatal parasitic disease in which effective immunity requires sufficient inflammation to control parasites while limiting immune-mediated tissue damage. Transforming growth factor-beta (TGFβ) is an important regulator of immune homeostasis and has been implicated in VL, but how it directly controls parasite-specific CD4⁺ T cell responses remains poorly understood. We used complementary transgenic mouse models with either enhanced or ablated TGFβ signalling in T cells during *Leishmania donovani* infection, combined with adoptive co-transfer of parasite-specific CD4⁺ T cells to distinguish cell-intrinsic effects. Enhanced TGFβ signalling impaired hepatic parasite control and suppressed CD4⁺ T cell immunity, reducing T helper 1 (Th1) cell differentiation, proliferation, accumulation of antigen-experienced cells, and expression of cytolytic molecules. Conversely, ablation of TGFβ signalling improved parasite control and promoted CD4⁺ T cell expansion and Th1 cell differentiation, while increasing expression of cytolytic molecules and reducing interleukin-10-producing type 1 regulatory T (Tr1) cells. Adoptive co-transfer experiments confirmed that TGFβ directly restrained the expansion and Th1 cell differentiation of parasite-specific CD4⁺ T cells and their acquisition of cytolytic features. Loss of signalling also impaired development of Tr1 cells and reduced expression of several chemokine receptors and co-inhibitory molecules associated with their regulatory function. However, enhanced signalling did not increase Tr1 cell development, indicating that the relationship between TGFβ signalling and immune regulation is not linear. TGFβ is a key cell-intrinsic regulator of CD4⁺ T cell fate during experimental VL. Rather than acting solely as a general suppressor of inflammation, it calibrates the balance between protective and regulatory immunity by controlling CD4⁺ T cell expansion, differentiation and effector function.

**Author summary:** Visceral leishmaniasis (VL) is a potentially fatal disease caused by *Leishmania* parasites. The immune system must generate a strong enough response to control these parasites while preventing excessive inflammation that can damage tissues. We investigated how transforming growth factor-beta (TGFβ), an important regulator of immune responses, helps maintain this balance. Using mice in which signalling by TGFβ was either increased or removed specifically in T cells, we found that this pathway strongly influenced the development and function of CD4⁺ T cells during infection. Increasing signalling suppressed the expansion of these cells and their development into inflammatory cells associated with parasite control. In contrast, removing signalling enhanced these responses and improved early parasite control, but also reduced the development of regulatory T cells that can limit inflammation. By studying parasite-specific T cells directly, we showed that many of these effects resulted from TGFβ acting within the T cells themselves. Our findings show that TGFβ does more than simply suppress immunity during VL. It helps determine the balance between CD4⁺ T cell responses that control parasites and those that regulate inflammation, providing new insight into how immunity is shaped during chronic infection.

## Introduction

Visceral leishmaniasis (VL) is the most severe manifestation of infection with protozoan parasites of the genus *Leishmania* and remains one of the world’s deadliest neglected tropical diseases [1, 2]. Caused primarily by *Leishmania donovani* and *L. infantum*, VL is characterised by hepatosplenomegaly, pancytopenia, chronic inflammation, and profound immune dysregulation, and is frequently fatal if left untreated [3]. There are 50,000–90,000 cases reported annually, and VL accounts for most leishmaniasis-associated deaths and remains endemic across the Indian subcontinent, East Africa, South America, and parts of the Mediterranean Basin and Central Asia [4]. Significant progress has been made towards disease control, with elimination programmes reducing case numbers by more than 95% in some endemic regions [5]. Nevertheless, an estimated 350 million people remain at risk, and factors including population displacement, emergence in new geographic areas, disruption of public health programmes, and persistent parasite reservoirs such as post-kala-azar dermal leishmaniasis (PKDL) patients threaten long-term elimination efforts [6–8]. A deeper understanding of the host–parasite interactions that drive disease pathogenesis and immunity remains essential for the development of improved interventions and sustainable disease control.

Experimental VL in mice has been instrumental in defining the immunological mechanisms that govern parasite control and persistence. Although susceptible mouse strains do not develop fatal disease, they exhibit a striking organ-specific dichotomy that mirrors the spectrum of human infection [9, 10]. Following *L. donovani* infection, parasites establish an acute infection in the liver that is progressively controlled through the formation of granulomas, highly organised immune structures in which parasite-specific T cells and activated macrophages cooperate to eliminate infected cells. In contrast, the spleen develops a chronic, progressive infection characterised by persistent parasite burden, elevated regulatory cytokines, T cell dysfunction, splenomegaly, and extensive disruption of tissue architecture [9–15]. These divergent outcomes within the same host provide a unique experimental system for comparing protective and non-protective immunity under highly comparable conditions. Importantly, the resolving hepatic response shares many features with asymptomatic human infection, whereas the chronically infected spleen recapitulates key pathological features of clinical VL, including chronic inflammation, immune suppression, and tissue remodelling [10]. As a result, experimental VLs in mice offers a powerful platform for dissecting the cellular and molecular pathways that determine whether anti-parasitic immunity leads to durable parasite control or persistent disease.

Transforming growth factor-β (TGF-β) is a pleiotropic cytokine that plays a central role in maintaining immune homeostasis by limiting excessive inflammation and regulating T cell differentiation [16]. Following activation from its latent form, TGF-β signals primarily through SMAD-dependent pathways to modulate gene expression across multiple immune cell types [16–18]. In infectious diseases, TGF-β suppresses protective immune responses by inhibiting T cell proliferation, macrophage activation, and the production of inflammatory cytokines such as IFN-γ, thereby limiting pathogen clearance [19, 20]. TGF-β also promotes immune regulation through the induction and maintenance of regulatory T cells and, in combination with other cytokines, contributes to the differentiation of Th17 cells [21]. In VL, elevated TGF-β levels are associated with impaired anti-parasitic immunity in both humans and experimental models, and the parasite can actively exploit this pathway to enhance its survival [22]. Beyond its direct immunosuppressive effects, TGF-β influences T cell fate decisions, including the maintenance of stem-like T cell populations and the regulation of transcriptional programs associated with effector and memory differentiation [23]. Collectively, these diverse activities position TGF-β as a key regulator of the balance between protective immunity, immune pathology, and chronic infection.

Despite substantial evidence implicating TGF-β in the regulation of immunity during visceral leishmaniasis [22, 24–28], its direct effects on parasite-specific CD4^+^ T cell responses remain poorly defined. To address this knowledge gap, we employed two complementary transgenic mouse models with either enhanced or ablated T cell-intrinsic TGF-β signalling during experimental VL. By combining these models with adoptive transfer approaches, we sought to define how TGF-β shapes the magnitude, differentiation, proliferative capacity, and effector functions of CD4^+^ T cells, and to determine the extent to which TGF-β signalling intrinsically regulates the balance between protective Th1 cell-mediated immunity and immunoregulatory T cell responses during chronic infection.

## Methods

All reagents, resources and equipment used in this study are described in Table 1.

**Table1.**
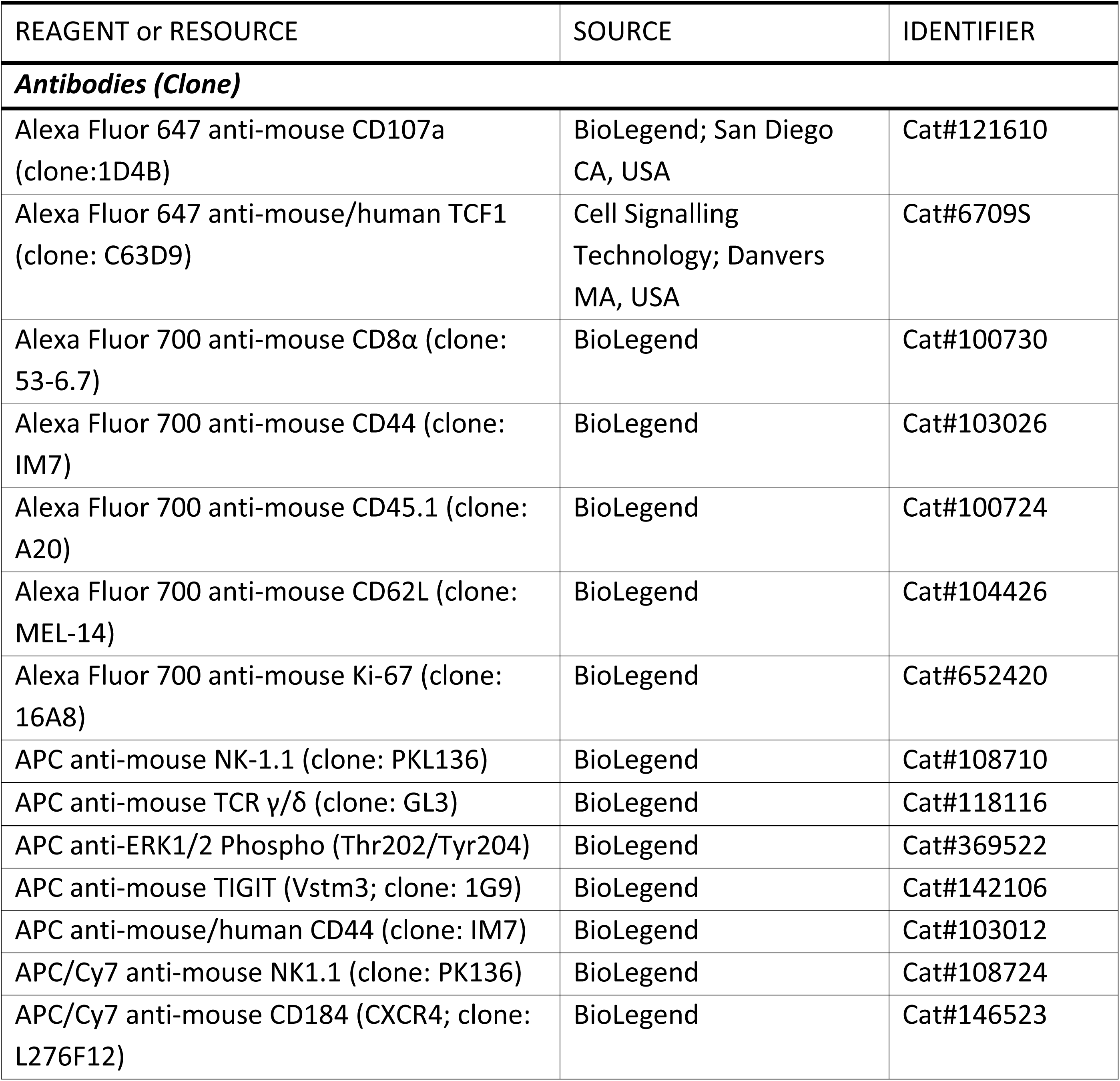
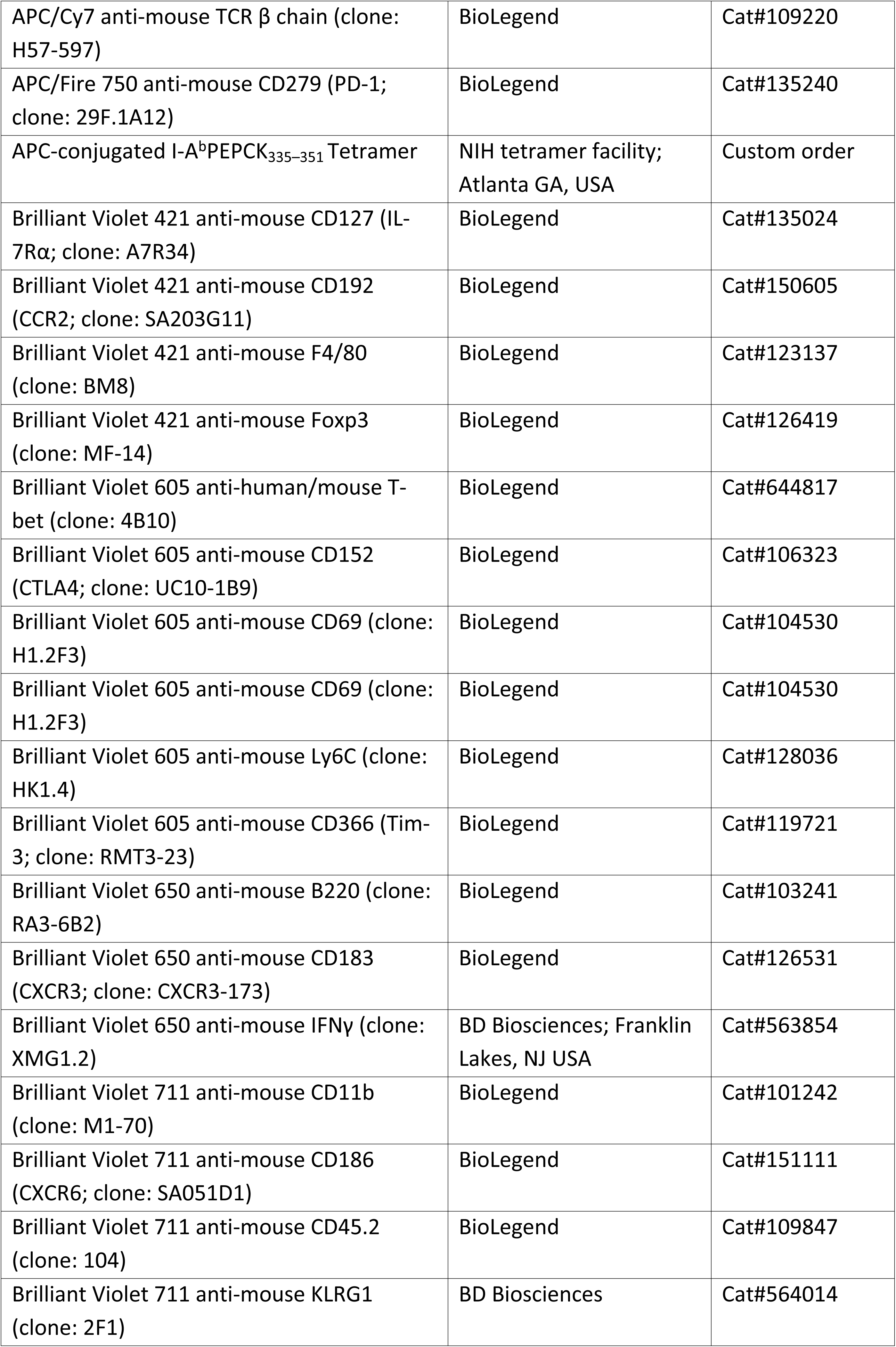
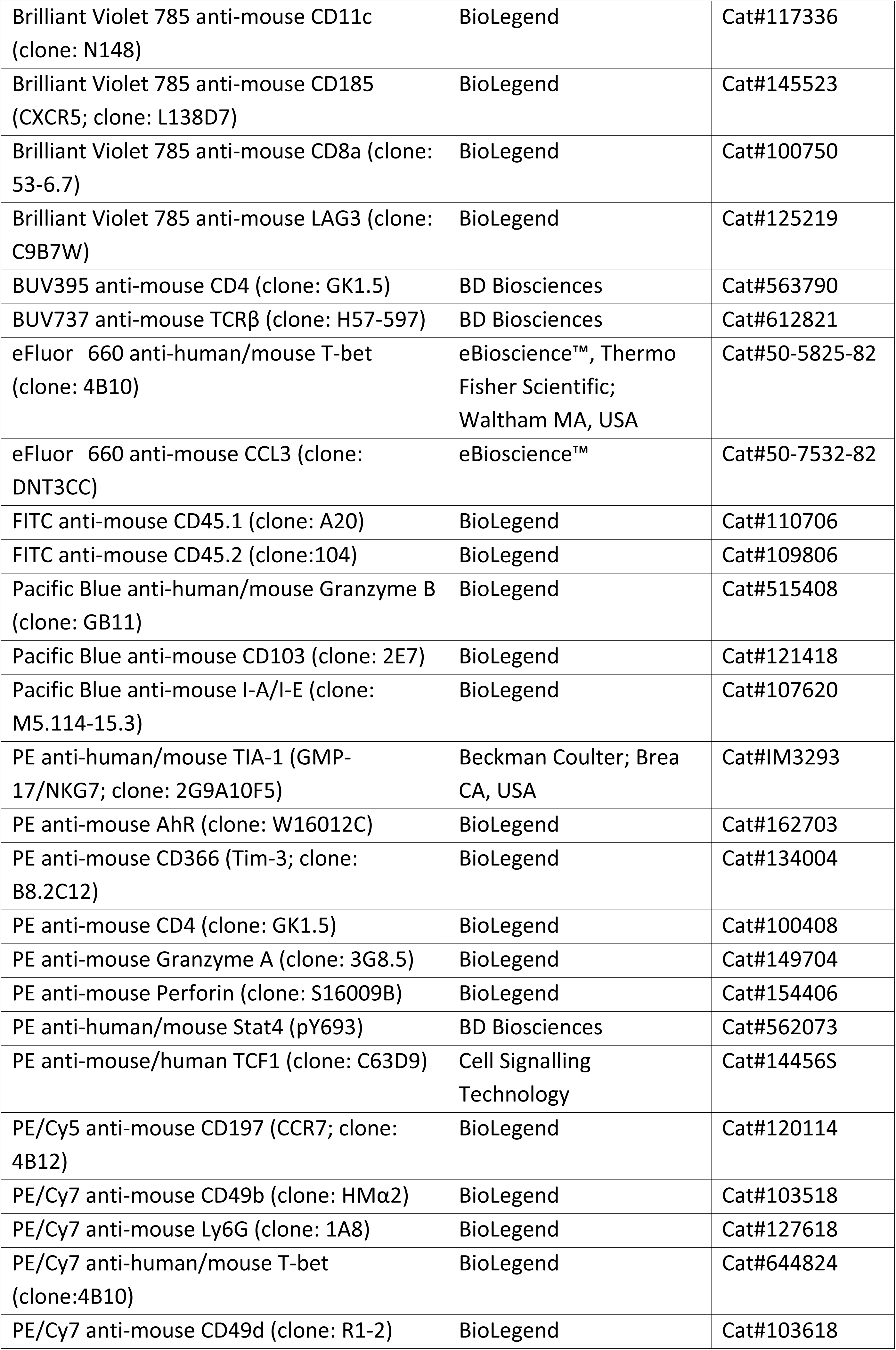
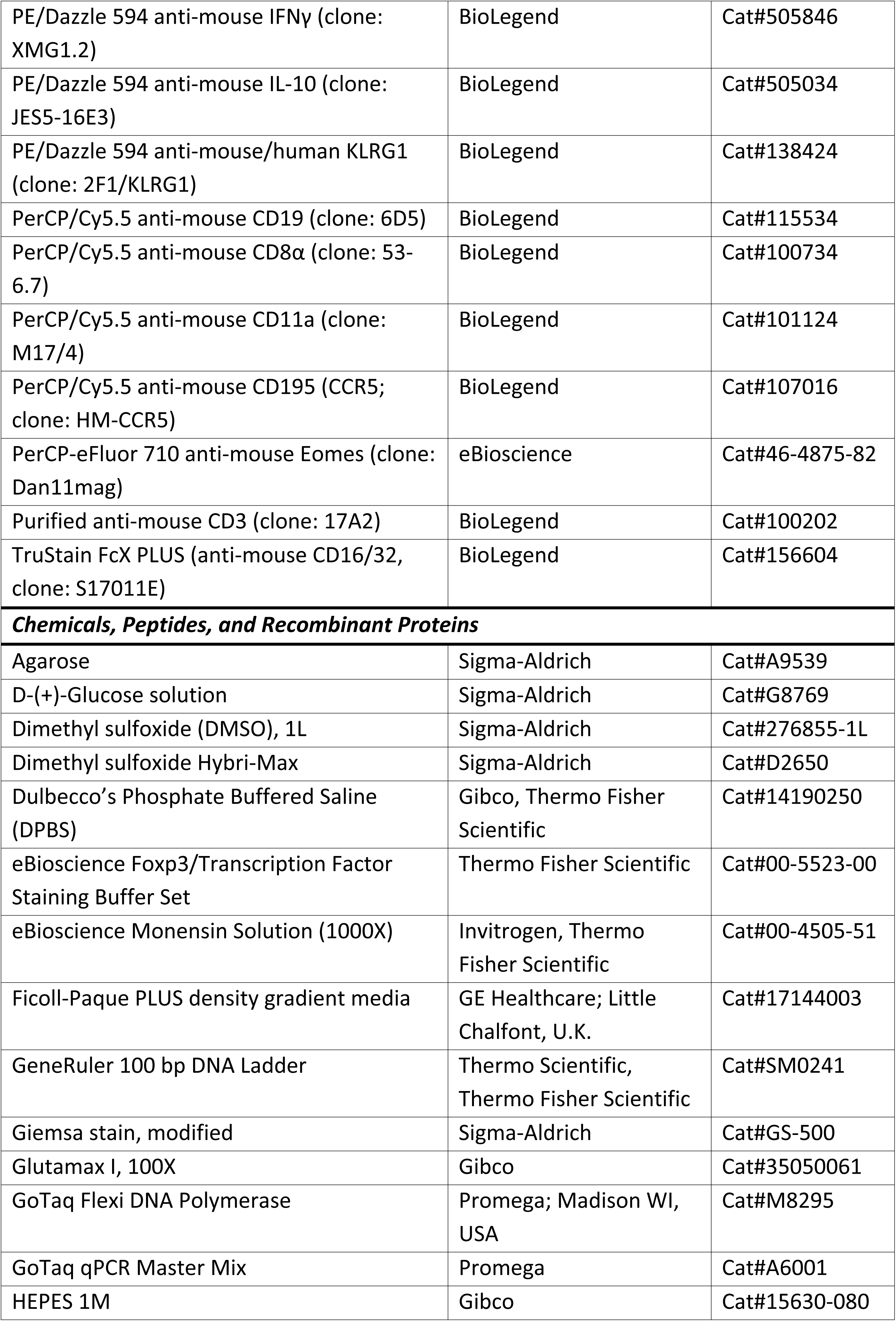
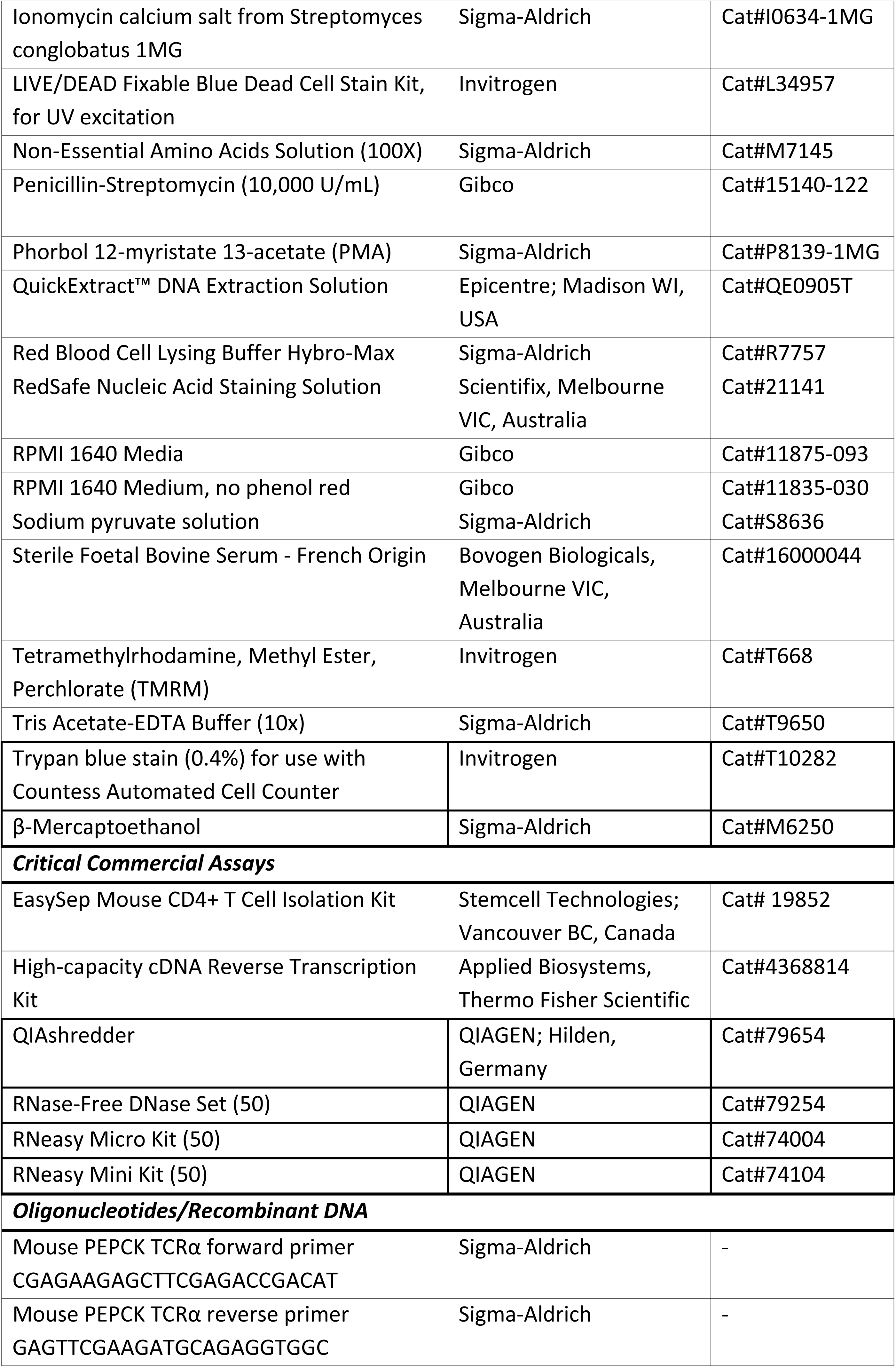
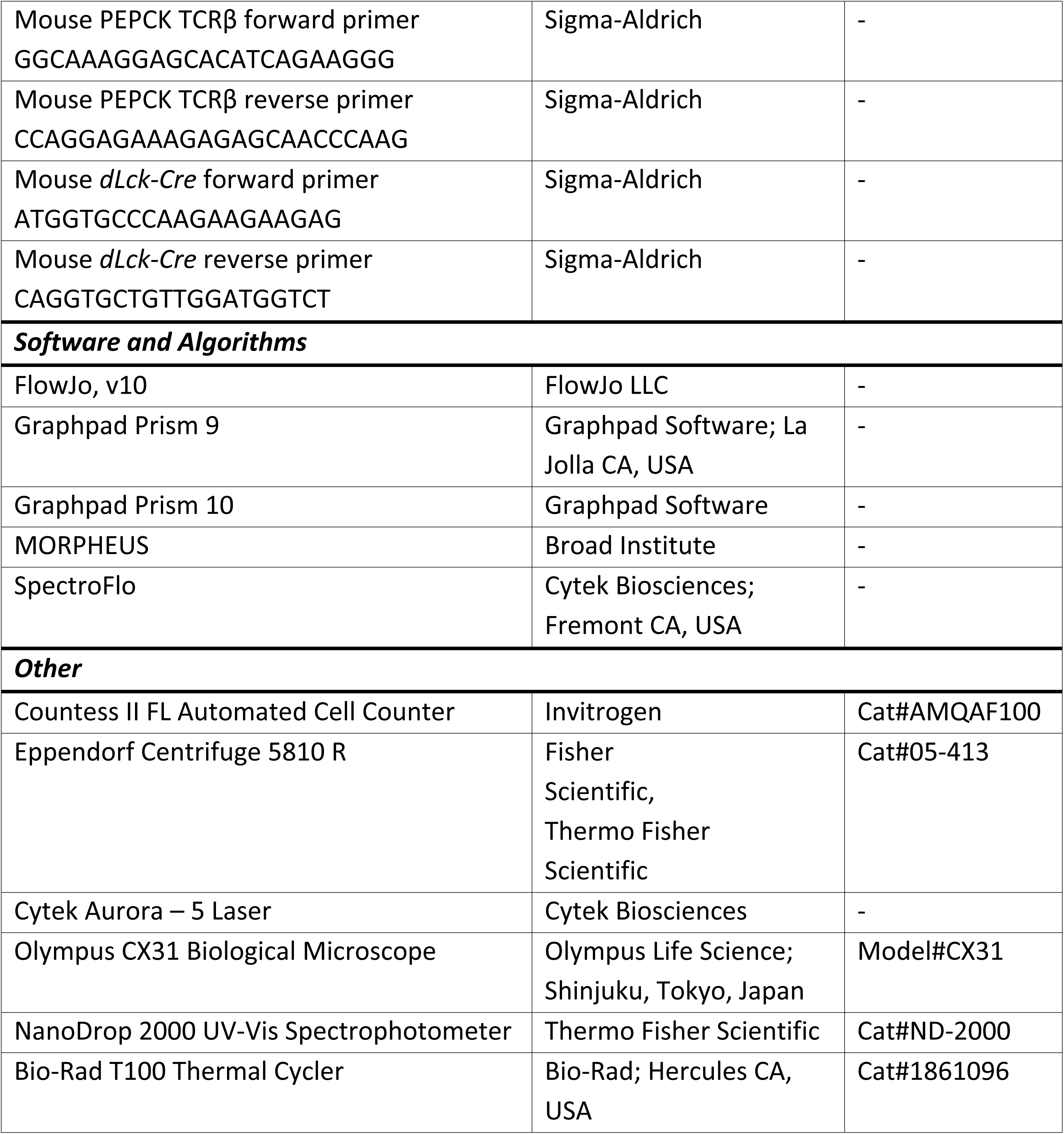
Reagents, Resources and Equipment.

### Mice

B6.CD45.1^+^ (B6.SJL-*Ptprc^a^* Pepc^b^/BoyJ) mice were purchased from Animal Resources Centre/ Ozgene (Canning Vale), B6.PEPCK [29] and B6.129S7-*Rag1^tm1Mom^*/J (*Rag1*^-/-^; JAX:002216) [30] were bred in house. C57BL/6 *lox*-stop-*lox*-*Tgfbr1*^CA^ (*Tgfbr1*^CA^) mice [31] and C57BL/6-129-*Tgfbr2^tm1karl^*/J (*Tgfbr2^fl/fl^*; JAX:012603) mice [32] were crossed with C57BL/6-Cg-Tg(*Lck-icre*)3779Nik/J (*dLck^cre^*; JAX:012837) mice [33] to produce mice with T cells either having constitutive TGFβR signalling (*Tgfbr1*^CA^) or lacking TGFβRII (B6.*Tgfbr2*^ΔT^), respectively. The Cre-negative littermates were used as controls. B6.PEPCK mice were crossed to *Ptprca* (CD45.1) mice to generate PEPCK x *CD45.1* (control, CD45.1^+^ CD45.2^+^), crossed to *Tgfbr1*^CA^ to generate T-cell specific TGFβ oversignalling mice (*Tgfbr1*^CA^ x PEPCK, CD45.2^+^), and *Tgfbr2*^ΔT^ to generate T-cell specific TGFβ signalling ablated mice (*Tgfbr2*^ΔT^ x PEPCK, CD45.2^+^). All mice were female, aged 8-12 weeks, and housed under pathogen-free conditions at the QIMR Berghofer Medical Research Institute Animal Facility (Herston QLD, Australia). Experimental mice use followed the “Australian Code of Practice for the Care and Use of Animals for Scientific Purposes” (Australian National Health and Medical Research Council) and was approved by the QIMR Berghofer Medical Research Institute Animal Ethics Committee (approval number: A1707-615M).

### Mouse genotyping

Mice requiring genotyping were ear notched, and the collected tissue was transferred into 40μL of QuickExtract DNA Extraction Solution (Epicentre; Madison WI, USA). DNA was extracted by heating the tissue sample in solution at 65°C for 6 minutes, then 98°C for 2 minutes. DNA primer oligos were custom ordered from Sigma-Aldrich (St. Louis MO, USA) (Table 1). Polymerase chain reactions (PCRs) performed using GoTaq Flexi DNA Polymerase (Promega; Madison WI, USA) were used to genotype all mice. PCRs were run on a Bio-Rad T100 Thermal Cycler (Bio-Rad Laboratories;17 Hercules CA, USA). Visualisation of PCR products used a 3% (w/v) agarose gel (Sigma-Aldrich) made up with 1x Tris Acetate-Ethylenediaminetetraacetic acid (EDTA) buffer (Sigma-Aldrich), containing RedSafe Nucleic Acid Staining Solution (Scientifix; Melbourne VIC, Australia) at 1:1000 dilution. PCR products were run in parallel with GeneRuler 100bp DNA ladder (Thermo Scientific, Waltham MA, USA) to visualise size.

### Leishmania donovani infections

*L. donovani* (LV9; MHOM/ET/67/HU3) amastigotes used for all infections were originally isolated from an Ethiopian patient in 1967 [34], and maintained through passage in B6.*Rag1*^-/-^ mice. Passage mice were euthanised via CO_2_ asphyxiation, and the spleen was removed and placed into 5mL of sterile Roswell Park Memorial Institute Medium 1640 (RPMI1640; Gibco, Thermo Fisher Scientific) supplemented with 100μg/mL penicillin-streptomycin (RPMI/PS; (PS; Gibco). A glass tissue grinder was used to homogenise the passage spleen and the resulting suspension was centrifuged in an Eppendorf Centrifuge 5810 R (Thermo Fisher Scientific) at 115*g* at room temperature (RT) for 5 minutes with brake off. The parasite-containing supernatant was extracted for further processing, and the cell debris pellet was discarded. The supernatant was centrifuged at 1960*g* for 15 minutes at RT, with brake-off. The supernatant was discarded, and the parasite pellet was incubated in 1mL of Red Blood Cell Lysis Buffer Hybri-Max (Sigma-Aldrich) for 5 minutes at RT, after which sterile RPMI/PS was added, and the parasites were centrifuged at 1960*g* for 15 minutes at RT. After the removal of the supernatant, the parasite pellet was resuspended in sterile RPMI/PS and the centrifugation step was repeated twice more for a total of three RPMI/PS washes post-RBC lysis. The pellet was resuspended in sterile RPMI/PS and the suspension was passed repeatedly through a 26G x ½” needle on a 1mL syringe (Terumo Medical; Somerset NJ, USA) to ensure homogenous parasite distribution in the suspension. 2μL were extracted from the suspension and loaded onto a Thoma cell counting chamber (Weber Scientific International; West Sussex, U.K.), and parasites were counted. The concentration of the parasite suspension was adjusted to 1×10^8^ parasites per mL using sterile RPMI/PS, and 200μL, or 2×10^7^ parasites, was administered per mice via tail vein intravenous (i.v.) injections, as previously described [35].

### Preparation of splenic single-cell suspensions

Mice were euthanised via CO_2_ asphyxiation, and the spleen was extracted via a mid-sagittal incision on the abdominal cavity. Spleens were weighed and collected into 1% (v/v) FBS (Bovogen Biologicals; Melbourne VIC, Australia) in PBS and were then mechanically passed through EASYstrainer 100μm cell strainer (Greiner Bio-One; Kremsmunster, Austria) using the back of a 5cc/mL syringe (Terumo Medical). The cell suspension was centrifuged in an Eppendorf Centrifuge 5810 R (Fisher Scientific) at 350*g* for 6 minutes at RT. The supernatant was removed, and the cell pellet was resuspended and incubated in 1mL of Red Blood Cell Lysis Buffer Hybri-Max (Sigma-Aldrich) for 6 minutes at RT. 1% (v/v) FBS in PBS was added to the suspension and the centrifuge step was repeated. Cells were resuspended in 1% (v/v) FBS in PBS and counted via dilutions in Dulbecco’s Phosphate Buffered Saline (DPBS) (1X) (Gibco™) and Trypan Blue Stain (Invitrogen, Thermo Fisher Scientific), using Countess Cell Counting Chamber Slides on the Countess II FL (both from Invitrogen), as instructed by the manufacturer.

### Preparation of hepatic single-cell suspensions

Mice were euthanised via CO_2_ asphyxiation, and a mid-sagittal incision was made on the abdominal cavity. Livers were perfused via the hepatic portal vein using 1x phosphate-buffered saline (PBS) before excision. Livers were weighed and collected into 1% (v/v) FBS (Bovogen Biologicals) in PBS and were then mechanically passed through a tea strainer fitted with a metal mesh using the back of a 10mL/cc syringe. The cell suspension was centrifuged in an Eppendorf Centrifuge 5810 R (Fisher Scientific) at 350*g* for 6 minutes at RT. The supernatant was removed, and the pellet was resuspended in 33% (v/v) Percoll Density Gradient Media (GE Healthcare; Little Chalfont, U.K.) and centrifuged at 575*g* for 14 minutes at RT with brake off. The supernatant containing the hepatocyte layer was removed, and the leucocyte pellet was resuspended and incubated in 1mL of Red Blood Cell Lysis Buffer Hybri-Max (Sigma-Aldrich) for 6 minutes at RT. 1% (v/v) FBS in PBS was added to the cell suspension and the resulting mix was centrifuged at 350*g* for 6 minutes at RT. The cell pellet was then resuspended in 1% (v/v) FBS in PBS and counted via dilutions in 1x DPBS (Gibco) and Trypan Blue Stain (Invitrogen), using Countess Cell Counting Chamber Slides on the Countess II FL (both from Invitrogen), as instructed by the manufacturer.

### Adoptive co-transfer of PEPCK CD4^+^ T cells

Naïve PEPCK mice were euthanised via CO_2_ asphyxiation, and the spleen was extracted via a mid-sagittal incision on the abdominal cavity. Spleens were processed into single-cell suspensions using sterile media and counted. Isolation of CD4^+^ T cells was conducted using EasySep Mouse CD4^+^ T Cell Isolation Kit (Stemcell technologies; Vancouver BC, Canada) according to manufacturer’s instructions, and isolated cells were resuspended in RPMI/PS. 100μL of isolated cells were stained with 1μg/mL of CD4-BUV395, TCRβ-BUV737, CD45.1-FITC and CD45.2-BV711 to assess post-isolation purity. Then, control PEPCK cells (CD45.1^+^, CD45.2^+^) were mixed with *Tgfbr1*^CA^ or *Tgfbr2*^ΔT^ PEPCK cells (both CD45.2^+^) at 1:1 and diluted to 2×10^5^ cells/mL using sterile RPMI/PS to create the injection mix. 200μL (4×10^4^) PEPCK cells were administered to each recipient *Ptprca* mouse (CD45.1^+^) via tail vein i.v. injections. At day 14 post-infection (p.i.), *Ptprca* mice were euthanised via CO_2_, and PEPCK cells of the spleen and liver were analysed to identify intrinsic differences caused by the gene modification of interest.

### Assessment of spleen and liver parasite burden

Sections of the spleen and liver were used to create impression smears, which were then stained in Giemsa (Sigma-Aldrich). *Leishmania* parasite burden measured in Leishman-Donovan Units (LDU) was quantified by the number of amastigotes per 1000 host nuclei multiplied by the organ weight (g). Slides were counted under x1000 magnification using a light microscope (Olympus CX31; Olympus Life Science, Shinjuku, Tokyo, Japan).

### Flow cytometry

All flow cytometric staining was carried out in Falcon 96-Well Clear Round Bottom Tissue Culture (TC)-Treated Cell Culture Microplates (Corning; Corning NY, USA), and all incubation steps were carried out in the dark to minimise fluorophore degradation. Splenic and hepatic cell suspensions were transferred into labelled wells and centrifuged at 575*g* for 2 minutes at RT. Supernatants were removed, and the pellet was resuspended in 30μL of staining cocktail made up using PBS containing TruStain FcX PLUS (BioLegend; San Diego CA, USA) and LIVE/DEAD Fixable Blue Dead Cell Stain (Invitrogen). Cells were incubated at 37°C for 15 minutes, after which cells were washed using PBS. Cells were then resuspended in 30μL of staining cocktail made using PBS containing surface staining antibodies (Table 1) relevant to the experiment and then incubated at 37°C for 20 minutes, followed by a PBS wash. After removal of the supernatant, cells were incubated in 100μL of fixation buffer from the eBioscience™ Foxp3/Transcription Factor Staining Buffer Set (Thermo Fisher Scientific) for 20 minutes at RT. Samples were then washed twice using the wash buffer from the same kit, at the previous centrifuge settings. Samples were resuspended in 30μL of intracellular staining mix made up using wash buffer, containing intracellular staining antibodies (Table 1) relevant to the experiment, and incubated at 37°C for 45 minutes. Samples were washed twice using the same wash buffer, resuspended in 100μL of PBS, and stored at 4°C before analysis. Samples were acquired on a Cytek Aurora (Cytek Biosciences; Fremont CA, USA) using SpectroFlo software (Cytek Biosciences), and data was analysed using FlowJo v10 (FlowJo, LLC; Ashland OR, USA). GraphPad Prism 9/10 (GraphPad Software; La Jolla CA, USA) was used for graphing and statistical analysis of flow cytometric data.

### Statistical analysis

For statistical comparisons made between mouse strains, and wild-type and TGFβR mutant cells, a Mann-Whitney test was used. Data was analysed using Graphpad Prism software (Graphpad Software; La Jolla CA, USA).

## Results

### TGF-β signalling to CD4^+^ T cells determines the outcome of L. donovani infection

To better understand the role of T cell-intrinsic TGF-β signalling during experimental VL, we employed two complementary transgenic mouse models that either enhanced or ablated TGF-β signalling in T cells (Figure 1A). The first is a gain-of-function model where *Tgfbr1*^CA^ mice exhibit constitutively active TGFβR1 [31] by T cells. The second is T cell-specific TGFβRII knockout (*Tgfbr2*^ΔT^) [32]. Following infection with *L. donovani*, these models revealed a critical role for T cell TGF-β signalling on parasite control in the first 14 days of infection, whereby constitutive TGF-β signalling limited control of parasite growth, albeit only in the liver, while lack of this pathway resulted in better parasite control, relative to control mice (Figure 1B-C).

**Figure 1.**
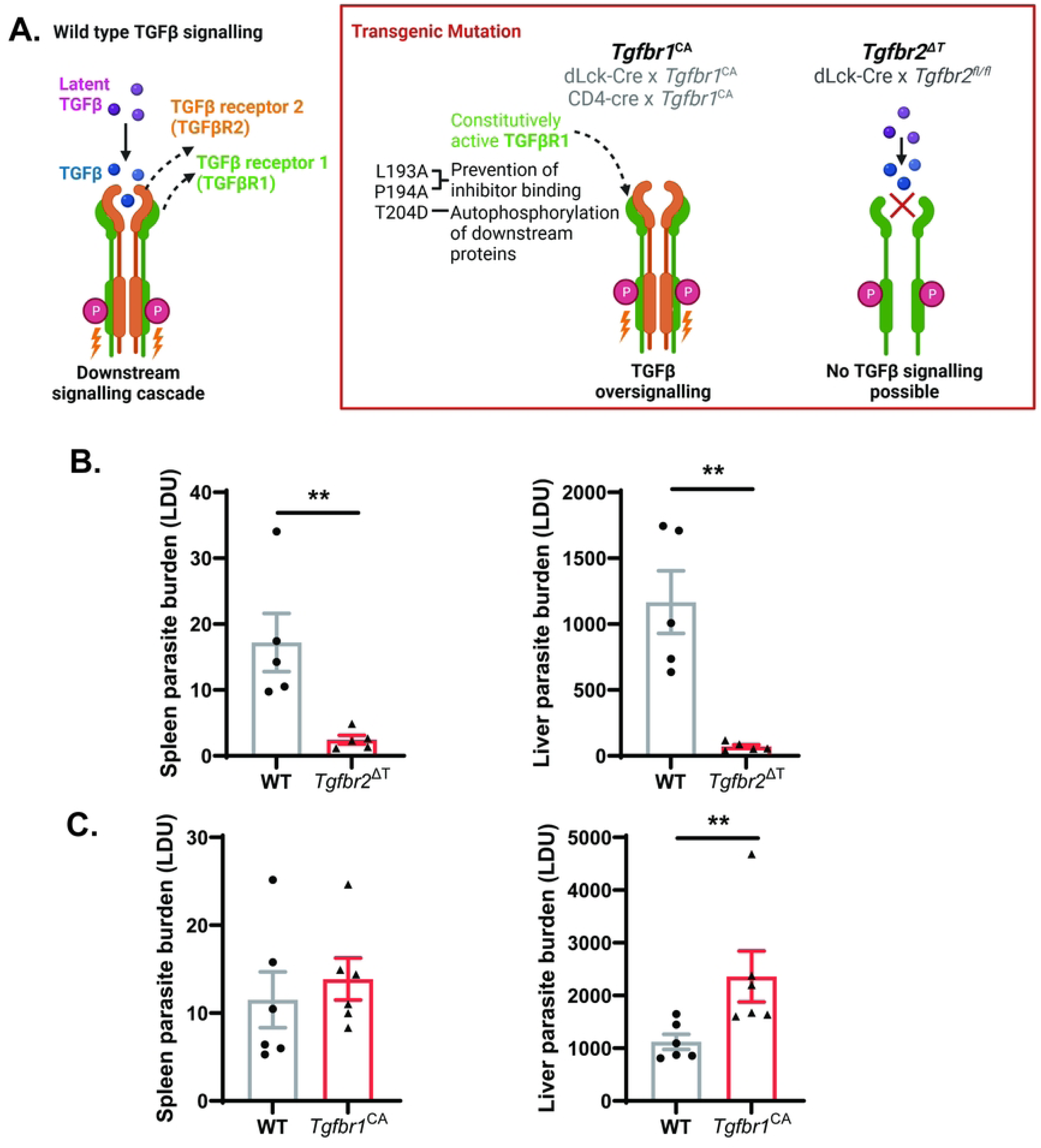
TGFβ signalling to T cells promotes *Leishmania donovani* growth. (**A**) TGFβ signalling via the TGFβ receptor (TGFβR) is depicted, as well as modifications to this pathway in transgenic mouse models with enhanced (*Tgfbr1*^CA^) or ablated (*Tgfbr2*^ΔT^) TGFβ signalling in T cells. Created in BioRender. Engwerda, C. (2026) BioRender.com/4e4b6kl. Parasite burdens in the spleen and liver, as indicated, of *Tgfbr2*^ΔT^ (**B**) or *Tgfbr1*^CA^ (**C**) mice, relative to wild-type (WT) controls at day 14 post-infection. A Mann-Whitney test was used to assess statistical significance, where ** p < 0.01, n = 5 per group (**B**) and 6 per group (**C**). Error bars indicate mean ± SEM.

### TGF-β over-signalling suppresses CD4⁺ T cell expansion, Th1 cell differentiation, and cytolytic functions during experimental VL

To determine how enhanced T cell-intrinsic TGF-β signalling influences developing CD4⁺ T cell responses during experimental VL, control (*Tgfbr1*^CA^ ^fl/fl^) and *Tgfbr1*^CA^ (*dLck-Cre*^+^ *Tgfbr1*^CA^) mice were infected with *L. donovani* and analysed 14 days later (Figure 2A). Total CD4⁺ T cell numbers were unchanged in the spleen but were significantly reduced in the liver of *Tgfbr1*^CA^ mice, suggesting that TGF-β over-signalling limits CD4⁺ T cell accumulation in highly inflammatory environments (Figure 2B). Consistent with a suppressive effect on effector differentiation, IFN-γ producing CD4^+^ T (Th1) cell frequencies were significantly reduced in both the spleen and liver of *Tgfbr1*^CA^ mice compared with controls (Figure 2C). IL-10-producing Th1 cells (type 1 regulatory (Tr) cells) are the major regulatory CD4^+^ T cells subset in experimental VL [36, 37], and frequencies of these cells were also reduced, although this effect was restricted to the spleen. In addition, CD4⁺ T cells from *Tgfbr1*^CA^ mice displayed significantly lower expression of the cytolytic and granule-associated proteins NKG7, CD107a, granzyme A and B, indicating that TGF-β signalling suppresses the acquisition of cytotoxic effector functions (Figure 2D).

**Figure 2.**
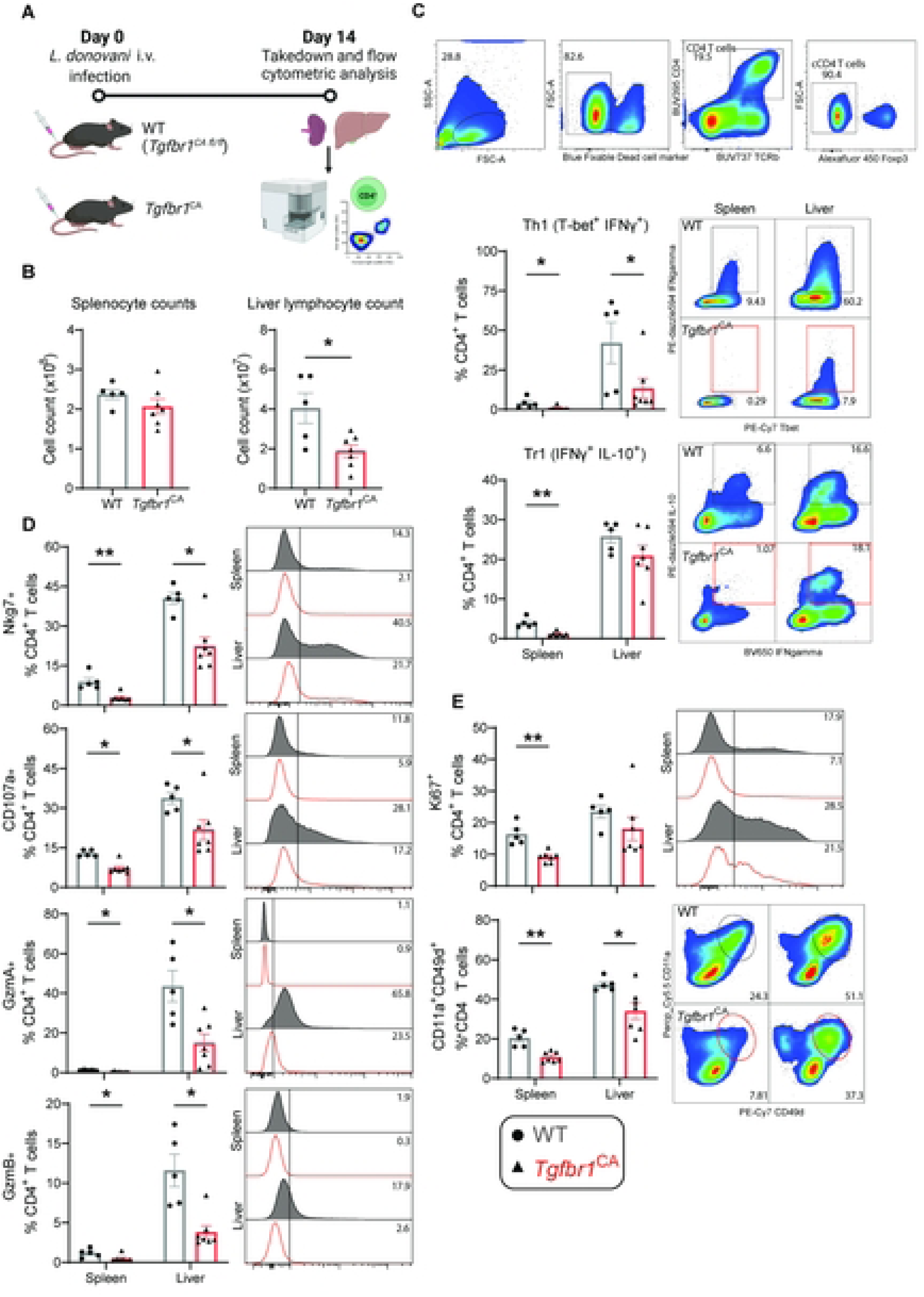
T cell specific TGFβ over-signalling results in suppression of T helper 1 cell responses during experimental visceral leishmaniasis. (**A**) Schematic of experimental outline for *Tgfbr1^CA^ ^fl/fl^* (WT) and *dLck-Cre x Tgfbr1^CA^ ^fl/fl^* (*Tgfbr1^CA^*) mice infected with *Leishmania donovani* and assessed at day 14 post-infection (p.i.). Created in BioRender. Engwerda, C. (2026) BioRender.com/6ot22yn. (**B**) Splenocyte and liver lymphocyte counts at day 14 post-infection. (**C**) Frequencies of Th1 (T-bet^+^ IFNγ^+^) and Tr1 (IFNγ^+^ IL-10^+^) cells within CD4^+^ T cells. (**D**) Expression of cytolytic molecules by CD4^+^ T cells. (**E**) Expression of the proliferation marker Ki67 and frequency of antigen specific CD4^+^ T cells marked by the co-expression of CD11a and CD49d. A Mann-Whitney test was used to assess statistical significance, where * p < 0.05 and ** p < 0.01, n = 5-7 mice per group, error bars indicate mean ± SEM.

We next assessed whether reduced CD4⁺ T cell accumulation reflected impaired proliferation and/or expansion. Expression of the proliferation marker Ki67 was significantly reduced in splenic CD4⁺ T cells from *Tgfbr1*^CA^ mice and showed a similar trend in the liver, although this wasn’t statistically significant (Figure 2E). Furthermore, the frequency of antigen-experienced CD4⁺ T cells, identified by co-expression of CD11a and CD49d [38], was significantly reduced in both organs. Although this decrease could reflect defects in activation, proliferation, or trafficking, the concomitant reduction in Ki67 expression suggests impaired proliferation is an important contributing factor. Collectively, these data demonstrate that enhanced TGF-β signalling broadly suppresses CD4⁺ T cell immunity during experimental VL, limiting Th1 cell differentiation, cytolytic programming, and the expansion of antigen-experienced cells.

We next used adoptive co-transfer experiments to determine whether these effects were mediated through cell-intrinsic mechanisms. PEPCK T cell receptor transgenic CD4^+^ T cells, which recognise a key antigen found in all disease-causing *Leishmania* species [29], were crossed with *Tgfbr1*^CA^ mice to produce parasite-specific CD4^+^ T cells with constitutively active TGFβR1 (*Tgfbr1*^CA^ PEPCK cells) and were homozygous for the CD45.2 allele. We also crossed PEPCK mice with congenic (CD45.1) C57BL/6 mice that were heterozygous for CD45 alleles (CD45.1/2). We then isolated splenic PEPCK cells from each mouse line and co-transferred them into congenic C57BL/6 mice that were homozygous for CD45.1, one day prior to infection with *L. donovani* (Figure 3A). The transferred cells were examined 14 days later by flow cytometry. Consistent with the previous observation that TGFβ has an inhibitory effect on the expansion of CD4^+^ T cells, we also found a significant reduction in the frequency of *Tgfbr1*^CA^ PEPCK cells relative to their WT control counterparts in both the spleen and liver (Figure 3B). Furthermore, there was a significant reduction in frequencies of Th1 cells in *Tgfbr1*^CA^ PEPCK cells relative to their WT controls (Figure 3C), but no change in Tr1 cell frequencies (Figure 3D). The very low frequencies of *Tgfbr1*^CA^ PEPCK cells meant we were unable to assess the impact of TGFβ over-signalling on activation and cytolytic markers.

**Figure 3.**
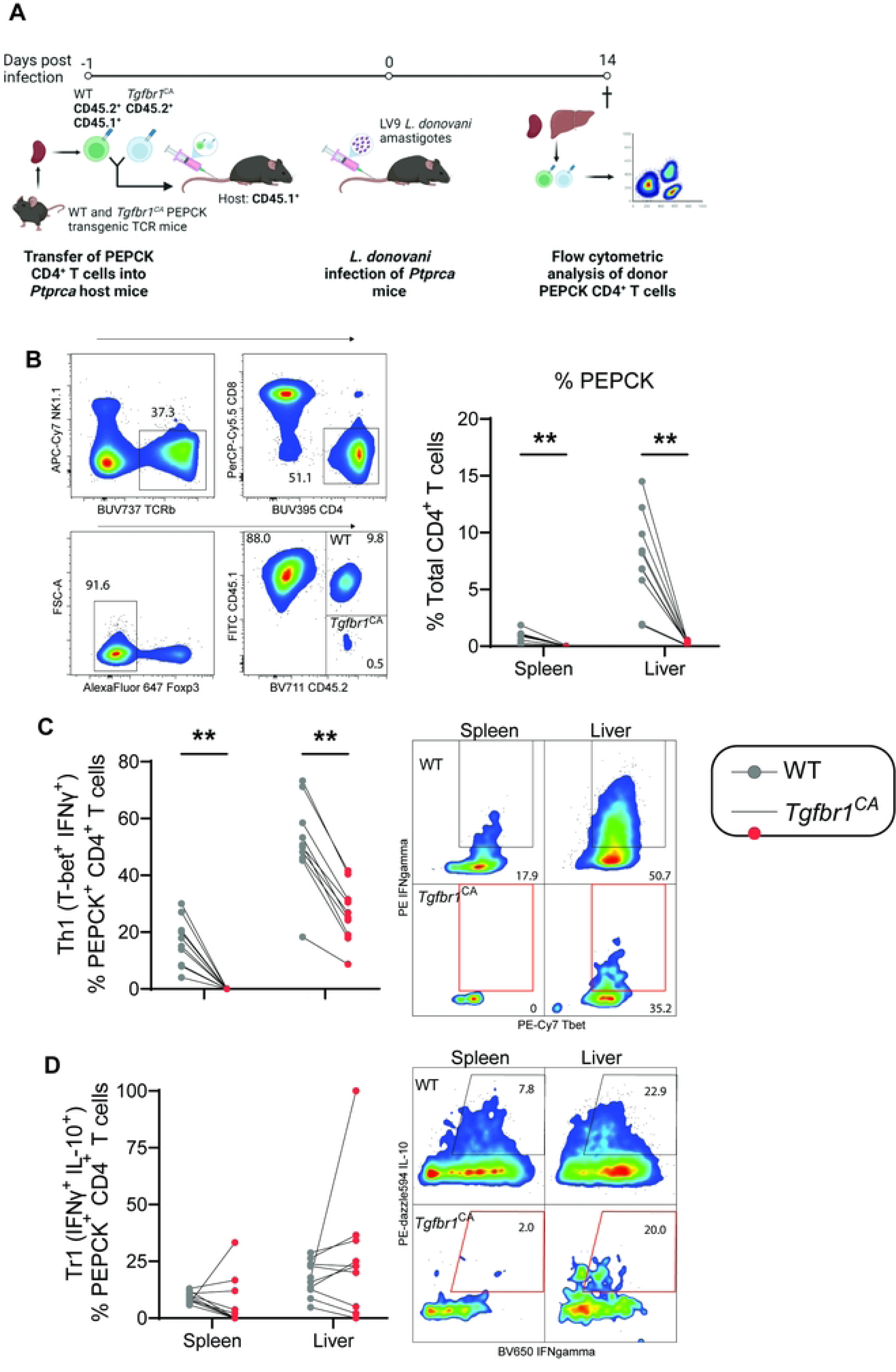
T cell-intrinsic TGFβ over-signalling suppresses expansion and differentiation of antigen-specific CD4^+^ T cells. (**A**) Schematic of experimental outline for *Tgfbr1*^CA^ ^fl/fl^ (WT) and *dLck-Cre* x *Tgfbr1*^CA^ ^fl/fl^ (*Tgfbr1*^CA^) PEPCK mice were transferred into recipient *Ptprca* (cd45.1) mice one day prior to *Leishmania donovani* infection and then assessed at day 14 post-infection. Created in BioRender. Engwerda, C. (2026) BioRender.com/6ot22yn. (**B**) Frequencies of WT and *Tgfbr1^CA^* PEPCK cells in the spleen and liver of the host mice. (**C**) Frequencies of Th1 (T-bet^+^ IFNγ^+^) and (**D**) Tr1 (IFNγ^+^ IL-10^+^) PEPCK cells in the spleen and liver. A Mann-Whitney test was used to assess statistical significance, where ** p < 0.01, n = 10 mice per group.

### Ablation of TGF-β signalling enhances Th1 differentiation, cytolytic programming, and expansion of CD4⁺ T cells during experimental VL

To determine how loss of T cell-intrinsic TGF-β signalling affects CD4⁺ T cell responses during experimental VL, WT (*Tgfbr2*^fl/fl^) and *Tgfbr2*^ΔT^ (*dLck-Cre*+ *Tgfbr2*^fl/fl^) mice were infected with *L. donovani* and analysed 14 days later (Figure 4A). In contrast to *Tgfbr1*^CA^ mice, *Tgfbr2*^ΔT^ mice exhibited significantly increased lymphocyte numbers in both the spleen and liver following infection, although CD4^+^ T cell numbers were already higher in the livers of *Tgfbr2*^ΔT^ mice, relative to controls, prior to infection (Figure 4B). This was accompanied by increased frequencies of Th1 cells and reduced frequencies of Tr1 cells, resulting in a marked shift towards a pro-inflammatory immune response (Figure 4C).

**Figure 4.**
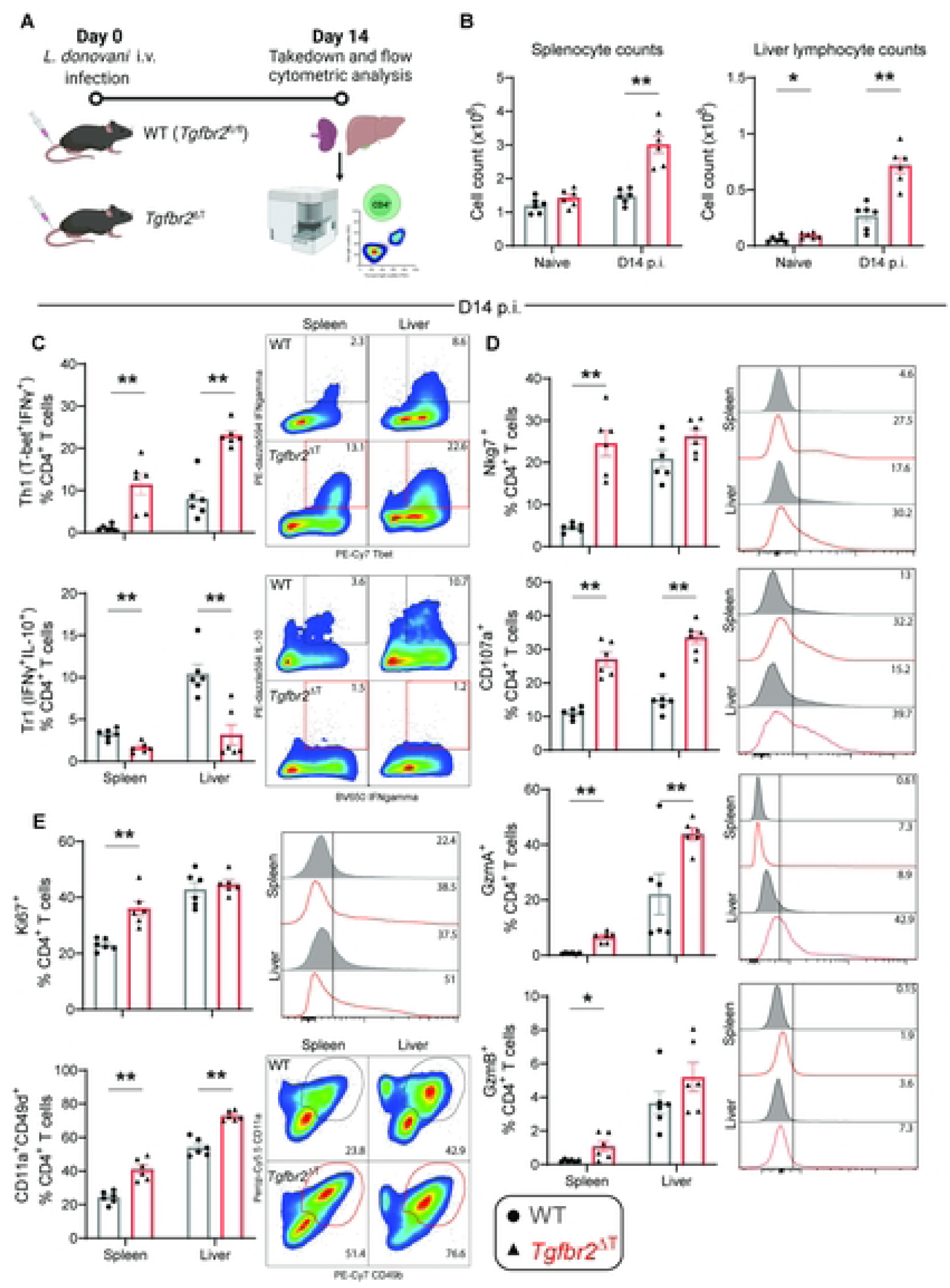
Ablation of TGFβ signalling results in enhanced Th1 cell responses during experimental visceral leishmaniasis. (**A**) Schematic of experimental outline for *Tgfbr2^fl/fl^*(WT) and *dLck-Cre* x *Tgfbr2^fl/fl^* (*Tgfbr2^ΔT^*) mice infected with *Leishmania donovani* and then analysed at day 14 post-infection (p.i.). Created in BioRender. Engwerda, C. (2026) BioRender.com/6ot22yn. (**B**) Splenocyte and liver lymphocyte counts at day 14 p.i. (**C**) Frequencies of Th1 (T-bet^+^ IFNγ^+^) and Tr1 (IFNγ^+^ IL-10^+^) cells within CD4^+^ T cells. (**D**) Expression of cytotoxicity-associated molecules by CD4^+^ T cells. (**E**) Expression of the proliferation marker Ki67 by CD4^+^ T cells, and frequency of antigen specific CD4^+^ T cells marked by co-expression of CD11a and CD49d. A Mann-Whitney test was used to assess statistical significance, where * p < 0.05 and ** p < 0.01, n = 6 mice per group, error bars indicate mean ± SEM.

Loss of TGF-β signalling also enhanced cytolytic programming of CD4⁺ T cells. Expression of cytolytic markers were consistently increased in the spleen of *Tgfbr2*^ΔT^ mice, whereas in the liver only CD107a and granzyme A were significantly elevated relative to WT controls (Figure 4D). Consistent with enhanced expansion, Ki67 expression was significantly increased in splenic CD4⁺ T cells from *Tgfbr2*^ΔT^ mice, although this effect was not observed in the liver (Figure 4E). Frequencies of antigen-experienced CD4⁺ T cells, identified by CD11a and CD49d co-expression, were also significantly increased in both organs, indicating enhanced expansion of parasite-specific cells. Together, these findings demonstrate that TGF-β signalling acts as a major constraint on CD4⁺ T cell immunity during experimental VL. Ablation of TGF-β signalling promoted Th1 cell differentiation, reduced Tr1 cell expansion, enhanced cytolytic programming, and increased the accumulation of CD4⁺ T cells.

We next used adoptive co-transfer experiments to determine the extent to which these effects were mediated through cell-intrinsic mechanisms. PEPCK cells were crossed with *Tgfbr2*^ΔT^ mice to produce parasite-specific CD4^+^ T cells lacking TGFβRII (*Tgfbr2*^ΔT^ PEPCK cells) and were homozygous for the CD45.2 allele. We also employed PEPCK mice crossed with congenic (CD45.1) C57BL/6 mice that were heterozygous for CD45 alleles (CD45.1/2). We isolated splenic PEPCK cells from each mouse line and co-transferred them into congenic C57BL/6 mice that were homozygous for CD45.1, one day prior to infection with *L. donovani*, as before (Figure 5A). The transferred cells were examined 14 days later by flow cytometry (Figure 5B-D). Consistent with the previous observation that abolishing TGFβ signalling improved Th1 cell responses and reduced Tr1 cell frequencies, we found an increase in Th1 cell frequency and a decrease in Tr1 cell frequency (Figure 5B). In addition, we found increased NKG7 and granzyme A expression in the spleen and liver by *Tgfbr2*^ΔT^ PEPCK cells, relative to control PEPCK cells, as well as increased CD107a expression in the liver and Ki67 expression in both the liver and spleen (Figure 5C). We also observed reduced expression of chemokine receptors and co-inhibitory receptors associated with the functions of Tr1 cells, including CCD5, PD-1 (spleen only) and TIGIT, but not Tim3 on *Tgfbr2*^ΔT^ PEPCK cells, relative to control PEPCK cells (Figure 5D). Together, these results indicate that cell intrinsic TGFβ signalling ablation on CD4^+^ T cells enhances Th1 cell responses and suppresses Tr1 cell development and functional capacity.

**Figure 5.**
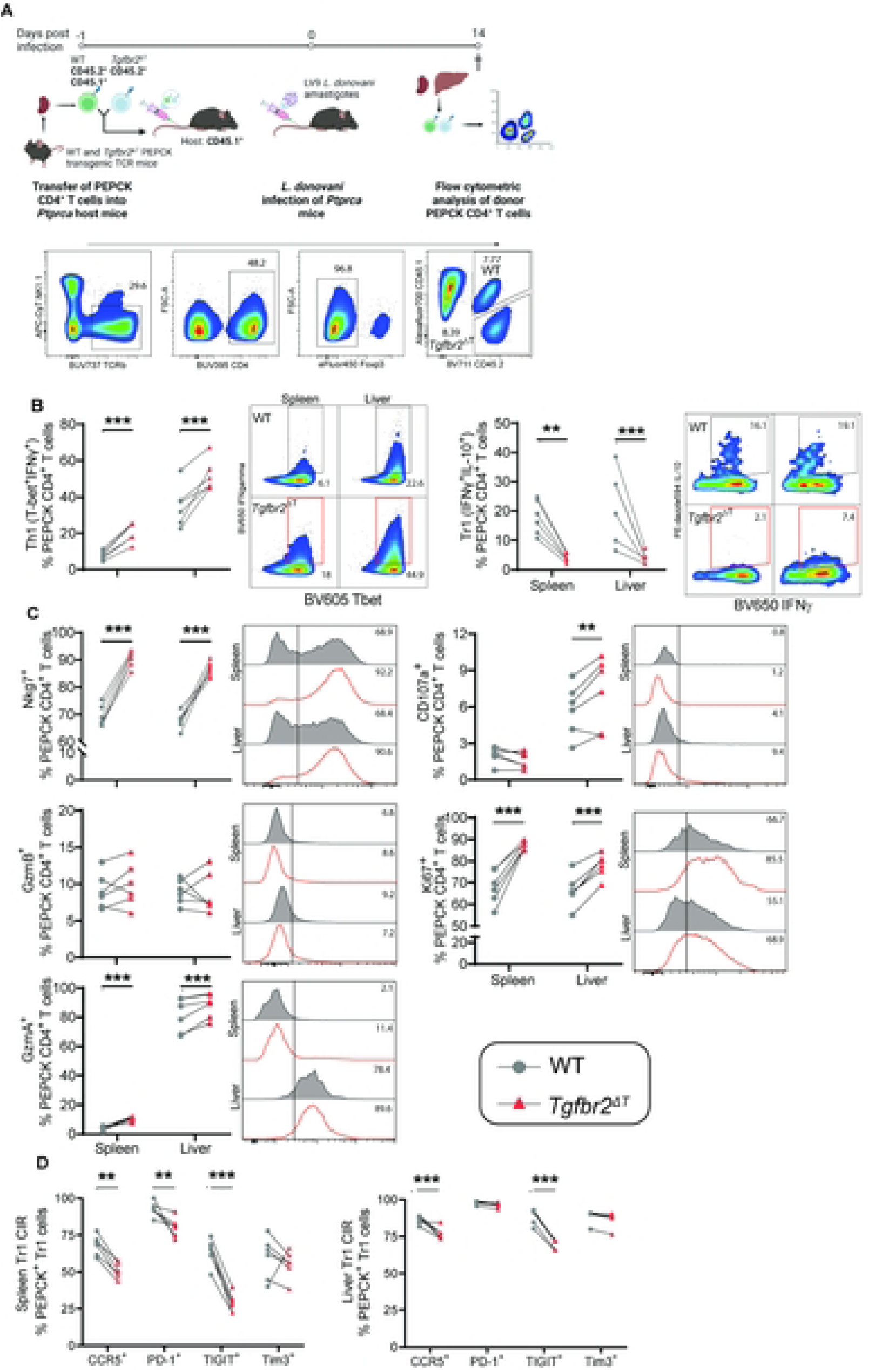
T cell-specific ablation of TGFβ signalling results in enhanced Th1 cell responses and reduced Tr1 cell responses during experimental visceral leishmaniasis. (**A**) Schematic of experimental outline for *Tgfbr2* ^fl/fl^ (WT) and *dLck-Cre* x *dLck-Cre* x *Tgfbr2^fl/fl^* (*Tgfbr2^ΔT^*) PEPCK cells transferred into recipient *Ptprca* (cd45.1) mice one day prior to *Leishmania donovani* infection then assessed and day 14 post-infection. Created in BioRender. Engwerda, C. (2026) BioRender.com/6ot22yn. (**B**) Frequencies of Th1 (T-bet^+^ IFNγ^+^) and Tr1 (IFNγ^+^ IL-10^+^) PEPCK cells. (**C**) Frequency of PEPCK CD4^+^ T cells expressing cytotoxicity-associated molecules and the proliferation marker Ki67. (**D**) Frequencies of PEPCK Tr1 cells expressing co-inhibitory receptors (CIRs) in the spleen and liver. A Mann-Whitney test was used to assess statistical significance, where ** p < 0.01 and *** p < 0.001, n = 5-6 mice per group, error bars indicate mean ± SEM.

## Discussion

In this study, we demonstrate that TGF-β is a key intrinsic regulator of CD4⁺ T cell immunity during experimental VL. Using complementary transgenic models with enhanced or ablated T cell-specific TGF-β signalling, we show that TGF-β constrains CD4⁺ T cell expansion, Th1 cell differentiation, and cytolytic programming, while also influencing the development and phenotype of regulatory Tr1 cells. The opposing phenotypes observed in *Tgfbr1*^CA^ and *Tgfbr2*^ΔT^ mice, together with adoptive co-transfer experiments, establish that many of these effects are mediated through cell-intrinsic TGF-β signalling pathways. Mechanistically, enhanced TGF-β signalling impaired the accumulation of antigen-experienced CD4⁺ T cells and suppressed their proliferative and effector capacity, whereas loss of TGF-β signalling promoted Th1 cell-driven inflammation, increased cytolytic marker expression, and enhanced expansion of parasite-specific CD4⁺ T cells. Interestingly, although absent TGF-β signalling impaired Tr1 cell development, excessive signalling didn’t cause enhanced Tr1 cell development, suggesting that either physiological levels of TGF-β are required for optimal Tr1 cell differentiation, while excessive signalling may become inhibitory or that reduced Th1 cell frequencies limited Tr1 cell development. Additionally, reduced expression of multiple co-inhibitory receptors on TGF-β-deficient Tr1 cells indicates that TGF-β contributes not only to Tr1 cell development but also to the maintenance of their regulatory phenotype.

Beyond its established role as an immunosuppressive cytokine, our findings suggest that TGF-β acts as a central regulator of CD4⁺ T cell fate during the establishment of chronic infection. TGF-β signalling influenced multiple aspects of the anti-parasitic response, including proliferation, effector differentiation, cytolytic programming, and regulatory cell development, highlighting its role as a key checkpoint balancing pathogen clearance and immune regulation. Consistent with previous reports that TGF-β suppresses T cell proliferation through inhibition of IL-2 signalling and cell cycle progression [25, 39–42], TGFβ over-signalling reduced Ki67 expression and the frequency of antigen-experienced CD4⁺ T cells. These defects likely reflect impaired proliferation together with reduced T cell activation, as TGF-β is known to inhibit multiple signalling events downstream of the T cell receptor [43–45]. Conversely, loss of TGF-β signalling enhanced CD4⁺ T cell expansion and increased the accumulation of antigen-experienced cells. Collectively, these findings indicate that TGF-β regulates not only the magnitude of the CD4⁺ T cell response but also its functional composition, extending previous studies focused on individual T cell subsets and positioning TGF-β as a broader coordinator of CD4⁺ T cell plasticity during chronic inflammatory conditions.

The identification of TGF-β signalling as a major intrinsic brake on anti-parasitic CD4⁺ T cell responses have potential therapeutic implications [19]. Host-directed immunotherapies that target inhibitory immune pathways, including TGF-β have shown considerable promise in cancer and chronic viral infections [19, 46, 47], and our data suggest that modulation of TGF-β signalling may represent a similar strategy for enhancing anti-parasitic immunity. While the pleiotropic functions of TGF-β would likely preclude prolonged systemic blockade [16, 20], transient, cell- or tissue-specific modulation of this pathway may provide a means to augment protective immunity in conjunction with anti-leishmanial chemotherapy or vaccination. Such approaches may be particularly relevant in settings of persistent infection, treatment failure, or impaired vaccine responsiveness.

An unexpected finding of this study was the profound effect of TGF-β signalling on the expression of cytolytic molecules by CD4⁺ T cells. Although cytolytic CD4⁺ T cells are increasingly recognised as important mediators of protection in chronic viral infections and cancer [48, 49], their contribution to immunity against *Leishmania* remains poorly understood. The increased expression of NKG7, granzymes, and CD107a following TGF-β signalling ablation suggests that cytolytic CD4⁺ T cells may represent an underappreciated effector population during VL. Future studies examining their capacity to directly kill infected host cells and their relationship to parasite control will be required to determine the functional significance of this population in anti-parasitic immunity.

Finally, our findings suggest that the relationship between TGF-β signalling and immune regulation is not linear. While loss of TGF-β signalling impaired Tr1 cell development and reduced expression of multiple co-inhibitory receptors, excessive TGF-β signalling didn’t cause expanded Tr1 cell frequencies. Although one explanation for this observation could be a reduced need for these cells because of diminished Th1 cell frequencies, these observations may also indicate that physiological levels of TGF-β signalling are required for optimal Tr1 cell differentiation and function. Such a model of TGF-β activity highlights the importance of signalling magnitude in determining T cell fate decisions and may help explain the diverse and sometimes contrasting effects attributed to TGF-β in chronic inflammatory diseases [50].

Taken together, our findings establish TGF-β as a critical determinant of CD4⁺ T cell fate during establishment of chronic *L. donovani* infection, acting not simply as a suppressor of inflammation but as a regulator of the balance between protective and regulatory immunity. By controlling the magnitude, differentiation, and functional capacity of parasite-specific CD4⁺ T cells, TGF-β shapes the quality of the host response to infection and influences the development of immune states associated with either parasite control or persistence. These results provide new mechanistic insight into how TGF-β signalling contributes to the immunological landscape of VL and highlight this pathway as a potential target for host-directed interventions aimed at enhancing anti-parasitic immunity while preserving immune homeostasis.

## Acknowledgements

We thank Dr Laura Mackay and Dr Axel Kallies (both from the Doherty Institute, University of Melbourne, Australia) for providing TGFβ receptor genetically modified mice used in this study to establish breeding colonies. We also thank Professor Jude Uzonna (University of Manitoba, Canada) for providing PEPCK T cell receptor transgenic mice to establish breeding colonies.

## References

1. Burza S, Croft SL, Boelaert M. Leishmaniasis. Lancet. 2018;392(10151):951-70. Epub 2018/08/22. doi: 10.1016/S0140-6736(18)31204-2. PubMed PMID: 30126638.

2. Pareyn M, Alves F, Burza S, Chakravarty J, Alvar J, Diro E, et al. Leishmaniasis. Nature Reviews Disease Primers. 2025;11(1):81. doi: 10.1038/s41572-025-00663-w.

3. Boelaert M, Sundar S. Leishmaniasis. In: Farrer J, Hotez PJ, Junghanss T, Kang G, Lalloo D, White N, editors. Manson’s Tropical Diseases. 23. Amsterdam: Elsevier; 2014. p. 631–51.

4. WHO. Leishmaniasis Fact Sheet 2023. Geneva: World Health Organization, 2023.

5. Sundar S. The story of elimination of visceral leishmaniasis (kala-azar) in India-Challenges towards sustainment. PLoS Negl Trop Dis. 2025;19(8):e0013321. Epub 2025/08/19. doi: 10.1371/journal.pntd.0013321. PubMed PMID: 40828749; PubMed Central PMCID: PMCPMC12364345.

6. Gedda MR, Singh B, Kumar D, Singh AK, Madhukar P, Upadhyay S, et al. Post kala-azar dermal leishmaniasis: A threat to elimination program. PLoS Negl Trop Dis. 2020;14(7):e0008221. Epub 2020/07/03. doi: 10.1371/journal.pntd.0008221. PubMed PMID: 32614818; PubMed Central PMCID: PMCPMC7332242.

7. Singh OP, Hasker E, Boelaert M, Sundar S. Elimination of visceral leishmaniasis on the Indian subcontinent. Lancet Infect Dis. 2016;16(12):e304–e9. Epub 2016/10/04. doi: 10.1016/S1473-3099(16)30140-2. PubMed PMID: 27692643; PubMed Central PMCID: PMCPMC5177523.

8. Singh OP, Tiwary P, Kushwaha AK, Singh SK, Singh DK, Lawyer P, et al. Xenodiagnosis to evaluate the infectiousness of humans to sandflies in an area endemic for visceral leishmaniasis in Bihar, India: a transmission-dynamics study. Lancet Microbe. 2021;2(1):e23–e31. Epub 2021/02/23. doi: 10.1016/s2666-5247(20)30166-x. PubMed PMID: 33615281; PubMed Central PMCID: PMCPMC7869864.

9. Engwerda CR, Kaye PM. Organ-specific immune responses associated with infectious disease. Immunology today. 2000;21(2):73–8. PubMed PMID: 10652464.

10. Engwerda CR, Kaye PM, Stäger S. Lymphocyte-mediated immune responses to Leishmania infection. Nat Rev Immunol. 2026. Epub 2026/04/23. doi: 10.1038/s41577-026-01294-2. PubMed PMID: 42020513.

11. Kaye PM, Svensson M, Ato M, Maroof A, Polley R, Stager S, et al. The immunopathology of experimental visceral leishmaniasis. Immunological reviews. 2004;201:239–53. doi: 10.1111/j.0105-2896.2004.00188.x. PubMed PMID: 15361245.

12. Stanley AC, Engwerda CR. Balancing immunity and pathology in visceral leishmaniasis. Immunology and cell biology. 2007;85(2):138–47. doi: 10.1038/sj.icb7100011. PubMed PMID: 17146466.

13. Engwerda C, Ato M, Cotterell S, Mynott T, Tschannerl A, Gorak-Stolinska P, et al. A Role for tumor necrosis factor-á in remodeling the splenic marginal zone during *Leishmania donovani* infection. The American Journal of Pathology. 2002;161(2):429–37. doi: 10.1016/s0002-9440(10)64199-5.

14. Engwerda CR, Ato M, Kaye PM. Macrophages, pathology and parasite persistence in experimental visceral leishmaniasis. Trends in parasitology. 2004;20(11):524–30. doi: 10.1016/j.pt.2004.08.009. PubMed PMID: 15471704.

15. Engwerda CR, Ato M, Stager S, Alexander CE, Stanley AC, Kaye PM. Distinct roles for lymphotoxin-alpha and tumor necrosis factor in the control of Leishmania donovani infection. The American journal of pathology. 2004;165(6):2123–33. PubMed PMID: 15579454; PubMed Central PMCID: PMC1618729.

16. Robertson IB, Rifkin DB. Regulation of the Bioavailability of TGF-β and TGF-β-Related Proteins. Cold Spring Harb Perspect Biol. 2016;8(6). Epub 2016/06/03. doi: 10.1101/cshperspect.a021907. PubMed PMID: 27252363; PubMed Central PMCID: PMCPMC4888822.

17. Cheifetz S, Hernandez H, Laiho M, ten Dijke P, Iwata KK, Massagué J. Distinct transforming growth factor-beta (TGF-beta) receptor subsets as determinants of cellular responsiveness to three TGF-beta isoforms. J Biol Chem. 1990;265(33):20533–8. Epub 1990/11/25. PubMed PMID: 1700790.

18. Shi Y, Massagué J. Mechanisms of TGF-beta signaling from cell membrane to the nucleus. Cell. 2003;113(6):685–700. Epub 2003/06/18. doi: 10.1016/s0092-8674(03)00432-x. PubMed PMID: 12809600.

19. Deng Z, Fan T, Xiao C, Tian H, Zheng Y, Li C, et al. TGF-β signaling in health, disease and therapeutics. Signal Transduction and Targeted Therapy. 2024;9(1):61. doi: 10.1038/s41392-024-01764-w.

20. Sanjabi S, Oh SA, Li MO. Regulation of the Immune Response by TGF-β: From Conception to Autoimmunity and Infection. Cold Spring Harb Perspect Biol. 2017;9(6). Epub 2017/01/22. doi: 10.1101/cshperspect.a022236. PubMed PMID: 28108486; PubMed Central PMCID: PMCPMC5453394.

21. Sanjabi S, Zenewicz LA, Kamanaka M, Flavell RA. Anti-inflammatory and pro-inflammatory roles of TGF-beta, IL-10, and IL-22 in immunity and autoimmunity. Curr Opin Pharmacol. 2009;9(4):447–53. Epub 2009/06/02. doi: 10.1016/j.coph.2009.04.008. PubMed PMID: 19481975; PubMed Central PMCID: PMCPMC2755239.

22. Barik S, Goswami S, Nanda PK, Sarkar A, Saha B, Sarkar A, et al. TGF-beta plays dual roles in immunity and pathogenesis in leishmaniasis. Cytokine. 2025;187:156865. doi: 10.1016/j.cyto.2025.156865.

23. Gabriel SS, Tsui C, Chisanga D, Weber F, Llano-León M, Gubser PM, et al. Transforming growth factor-beta-regulated mTOR activity preserves cellular metabolism to maintain long-term T cell responses in chronic infection. Immunity. 2021;54(8):1698–714.e5. doi: 10.1016/j.immuni.2021.06.007.

24. Elmekki MA, Elhassan MM, Ozbak HA, Mukhtar MM. Elevated TGF-beta levels in drug-resistant visceral leishmaniasis. Ann Saudi Med. 2016;36(1):73–7. Epub 2016/02/29. doi: 10.5144/0256-4947.2016.73. PubMed PMID: 26922691; PubMed Central PMCID: PMCPMC6074269 report related to this article.

25. Gorelik L, Constant S, Flavell RA. Mechanism of transforming growth factor beta-induced inhibition of T helper type 1 differentiation. J Exp Med. 2002;195(11):1499–505. Epub 2002/06/05. doi: 10.1084/jem.20012076. PubMed PMID: 12045248; PubMed Central PMCID: PMCPMC2193549.

26. Kupani M, Sharma S, Pandey RK, Kumar R, Sundar S, Mehrotra S. IL-10 and TGF-β Induced Arginase Expression Contributes to Deficient Nitric Oxide Response in Human Visceral Leishmaniasis. Front Cell Infect Microbiol. 2020;10:614165. Epub 2021/03/09. doi: 10.3389/fcimb.2020.614165. PubMed PMID: 33680983; PubMed Central PMCID: PMCPMC7930829.

27. Rodrigues OR, Marques C, Soares-Clemente M, Ferronha MH, Santos-Gomes GM. Identification of regulatory T cells during experimental Leishmania infantum infection. Immunobiology. 2009;214(2):101–11. Epub 2009/01/27. doi: 10.1016/j.imbio.2008.07.001. PubMed PMID: 19167988.

28. Wilson ME, Young BM, Davidson BL, Mente KA, McGowan SE. The importance of TGF-beta in murine visceral leishmaniasis. J Immunol. 1998;161(11):6148–55. Epub 1998/12/02. PubMed PMID: 9834100.

29. Barazandeh AF, Mou Z, Ikeogu N, Mejia EM, Edechi CA, Zhang WW, et al. The Phosphoenolpyruvate Carboxykinase Is a Key Metabolic Enzyme and Critical Virulence Factor of Leishmania major. J Immunol. 2021;206(5):1013–26. Epub 2021/01/20. doi: 10.4049/jimmunol.2000517. PubMed PMID: 33462138.

30. Mombaerts P, Iacomini J, Johnson RS, Herrup K, Tonegawa S, Papaioannou VE. RAG-1-deficient mice have no mature B and T lymphocytes. Cell. 1992;68(5):869–77. Epub 1992/03/06. doi: 10.1016/0092-8674(92)90030-g. PubMed PMID: 1547488.

31. Bartholin L, Cyprian FS, Vincent D, Garcia CN, Martel S, Horvat B, et al. Generation of mice with conditionally activated transforming growth factor beta signaling through the TbetaRI/ALK5 receptor. Genesis. 2008;46(12):724–31. Epub 2008/09/30. doi: 10.1002/dvg.20425. PubMed PMID: 18821589.

32. Levéen P, Larsson J, Ehinger M, Cilio CM, Sundler M, Sjöstrand LJ, et al. Induced disruption of the transforming growth factor beta type II receptor gene in mice causes a lethal inflammatory disorder that is transplantable. Blood. 2002;100(2):560–8. Epub 2002/07/02. doi: 10.1182/blood.v100.2.560. PubMed PMID: 12091349.

33. Wang Q, Strong J, Killeen N. Homeostatic competition among T cells revealed by conditional inactivation of the mouse Cd4 gene. J Exp Med. 2001;194(12):1721–30. Epub 2001/12/19. doi: 10.1084/jem.194.12.1721. PubMed PMID: 11748274; PubMed Central PMCID: PMCPMC2193581.

34. Bradley DJ, Kirkley J. Regulation of Leishmania populations within the host. I. the variable course of Leishmania donovani infections in mice. Clin Exp Immunol. 1977;30(1):119–29. Epub 1977/10/01. PubMed PMID: 606433; PubMed Central PMCID: PMCPMC1541173.

35. Ng SS, De Labastida Rivera F, Yan J, Corvino D, Das I, Zhang P, et al. The NK cell granule protein NKG7 regulates cytotoxic granule exocytosis and inflammation. Nat Immunol. 2020;21(10):1205–18. Epub 2020/08/26. doi: 10.1038/s41590-020-0758-6. PubMed PMID: 32839608; PubMed Central PMCID: PMCPMC7965849.

36. Bunn PT, Montes de Oca M, de Labastida Rivera F, Kumar R, Ng SS, Edwards CL, et al. Distinct Roles for CD4(+) Foxp3(+) Regulatory T Cells and IL-10-Mediated Immunoregulatory Mechanisms during Experimental Visceral Leishmaniasis Caused by Leishmania donovani. J Immunol. 2018;201(11):3362–72. Epub 2018/10/26. doi: 10.4049/jimmunol.1701582. PubMed PMID: 30355785.

37. Montes de Oca M, Kumar R, de Labastida Rivera F, Amante FH, Sheel M, Faleiro RJ, et al. Blimp-1-Dependent IL-10 Production by Tr1 Cells Regulates TNF-Mediated Tissue Pathology. PLoS pathogens. 2016;12(1):e1005398. doi: 10.1371/journal.ppat.1005398. PubMed PMID: 26765224.

38. McDermott DS, Varga SM. Quantifying antigen-specific CD4 T cells during a viral infection: CD4 T cell responses are larger than we think. J Immunol. 2011;187(11):5568–76. Epub 2011/11/02. doi: 10.4049/jimmunol.1102104. PubMed PMID: 22043009; PubMed Central PMCID: PMCPMC3221938.

39. Lucas PJ, Kim SJ, Melby SJ, Gress RE. Disruption of T cell homeostasis in mice expressing a T cell-specific dominant negative transforming growth factor beta II receptor. J Exp Med. 2000;191(7):1187–96. Epub 2000/04/05. doi: 10.1084/jem.191.7.1187. PubMed PMID: 10748236; PubMed Central PMCID: PMCPMC2193176.

40. McKarns SC, Schwartz RH, Kaminski NE. Smad3 is essential for TGF-beta 1 to suppress IL-2 production and TCR-induced proliferation, but not IL-2-induced proliferation. J Immunol. 2004;172(7):4275–84. Epub 2004/03/23. doi: 10.4049/jimmunol.172.7.4275. PubMed PMID: 15034041.

41. Ruegemer JJ, Ho SN, Augustine JA, Schlager JW, Bell MP, McKean DJ, et al. Regulatory effects of transforming growth factor-beta on IL-2- and IL-4-dependent T cell-cycle progression. J Immunol. 1990;144(5):1767–76. Epub 1990/03/01. PubMed PMID: 2407783.

42. Wolfraim LA, Walz TM, James Z, Fernandez T, Letterio JJ. p21Cip1 and p27Kip1 act in synergy to alter the sensitivity of naive T cells to TGF-beta-mediated G1 arrest through modulation of IL-2 responsiveness. J Immunol. 2004;173(5):3093–102. Epub 2004/08/24. doi: 10.4049/jimmunol.173.5.3093. PubMed PMID: 15322169.

43. Boussiotis VA, Chen ZM, Zeller JC, Murphy WJ, Berezovskaya A, Narula S, et al. Altered T-cell receptor + CD28-mediated signaling and blocked cell cycle progression in interleukin 10 and transforming growth factor-beta-treated alloreactive T cells that do not induce graft-versus-host disease. Blood. 2001;97(2):565–71. Epub 2001/01/12. doi: 10.1182/blood.v97.2.565. PubMed PMID: 11154238.

44. Chen CH, Seguin-Devaux C, Burke NA, Oriss TB, Watkins SC, Clipstone N, et al. Transforming growth factor beta blocks Tec kinase phosphorylation, Ca2+ influx, and NFATc translocation causing inhibition of T cell differentiation. J Exp Med. 2003;197(12):1689–99. Epub 2003/06/18. doi: 10.1084/jem.20021170. PubMed PMID: 12810687; PubMed Central PMCID: PMCPMC2193945.

45. Giroux M, Delisle JS, O’Brien A, Hébert MJ, Perreault C. T cell activation leads to protein kinase C theta-dependent inhibition of TGF-beta signaling. J Immunol. 2010;185(3):1568–76. Epub 2010/07/02. doi: 10.4049/jimmunol.1000137. PubMed PMID: 20592275.

46. Wallis RS, O’Garra A, Sher A, Wack A. Host-directed immunotherapy of viral and bacterial infections: past, present and future. Nat Rev Immunol. 2023;23(2):121–33. Epub 2022/06/08. doi: 10.1038/s41577-022-00734-z. PubMed PMID: 35672482; PubMed Central PMCID: PMCPMC9171745.

47. Wykes MN, Lewin SR. Immune checkpoint blockade in infectious diseases. Nat Rev Immunol. 2018;18(2):91–104. Epub 2017/10/11. doi: 10.1038/nri.2017.112. PubMed PMID: 28990586; PubMed Central PMCID: PMCPMC5991909.

48. Lai L, Ran S, Li Y, Cui J, Zhang X, Yu J, et al. Cytotoxic CD4+ T cells: origin, biological functions, diseases and therapeutic targets. Signal Transduction and Targeted Therapy. 2026;11(1):85. doi: 10.1038/s41392-025-02533-z.

49. Malyshkina A, Brüggemann A, Paschen A, Dittmer U. Cytotoxic CD4+ T cells in chronic viral infections and cancer. Frontiers in immunology. 2023;Volume 14 - 2023. doi: 10.3389/fimmu.2023.1271236.

50. Moreau JM, Velegraki M, Bolyard C, Rosenblum MD, Li Z. Transforming growth factor–β1 in regulatory T cell biology. Science Immunology. 2022;7(69):eabi4613. doi: doi:10.1126/sciimmunol.abi4613.

